# APOE, Immune Factors, Sex, and Diet Interact to Shape Brain Networks in Mouse Models of Aging

**DOI:** 10.1101/2023.10.04.560954

**Authors:** Steven Winter, Ali Mahzarnia, Robert J Anderson, Zay Yar Han, Jessica Tremblay, Jacques Stout, Hae Sol Moon, Daniel Marcellino, David B. Dunson, Alexandra Badea

## Abstract

Alzheimer’s disease (AD) presents complex challenges due to its multifactorial nature, poorly understood etiology, and late detection. The mechanisms through which genetic, fixed and modifiable risk factors influence susceptibility to AD are under intense investigation, yet the impact of unique risk factors on brain networks is difficult to disentangle, and their interactions remain unclear. To model multiple risk factors including APOE genotype, age, sex, diet, and immunity we leveraged mice expressing the human APOE and NOS2 genes, conferring a reduced immune response compared to mouse Nos2. Employing graph analyses of brain connectomes derived from accelerated diffusion-weighted MRI, we assessed the global and local impact of risk factors in the absence of AD pathology. Aging and a high-fat diet impacted extensive networks comprising AD-vulnerable regions, including the temporal association cortex, amygdala, and the periaqueductal gray, involved in stress responses. Sex impacted networks including sexually dimorphic regions (thalamus, insula, hypothalamus) and key memory-processing areas (fimbria, septum). APOE genotypes modulated connectivity in memory, sensory, and motor regions, while diet and immunity both impacted the insula and hypothalamus. Notably, these risk factors converged on a circuit comprising 63 of 54,946 total connections (0.11% of the connectome), highlighting shared vulnerability amongst multiple AD risk factors in regions essential for sensory integration, emotional regulation, decision making, motor coordination, memory, homeostasis, and interoception. These network-based biomarkers hold translational value for distinguishing high-risk versus low-risk participants at preclinical AD stages, suggest circuits as potential therapeutic targets, and advance our understanding of network fingerprints associated with AD risk.

**Significance Statement:** Current interventions for Alzheimer’s disease (AD) do not provide a cure, and are delivered years after neuropathological onset. Addressing the impact of risk factors on brain networks holds promises for early detection, prevention, and revealing putative therapeutic targets at preclinical stages. We utilized six mouse models to investigate the impact of factors, including APOE genotype, age, sex, immunity, and diet, on brain networks. Large structural connectomes were derived from high resolution compressed sensing diffusion MRI. A highly parallelized graph classification identified subnetworks associated with unique risk factors, revealing their network fingerprints, and a common network composed of 63 connections with shared vulnerability to all risk factors. APOE genotype specific immune signatures support the design of interventions tailored to risk profiles.

## Introduction

Late onset Alzheimer’s disease (LOAD) is a complex neurodegenerative disease affecting >13% of people over 75 [1]. In the absence of a cure, a better understanding of risk factors is central to early detection, prevention, and unlocking the potential of early interventions. Age is the largest LOAD risk factor. Amongst risk genes, APOE has the strongest impact [2, 3] [4], the APOE2 allele being thought of as protective, APOE3 as neutral, and APOE4 conferring the greatest risk [5, 6]. It is however unclear how APOE4 interacts with age, immunity [7], metabolic status [8], and sex to increase vulnerability; females constitute 2/3 of LOAD patients [9]. Modifiable risk factors include a history of brain injuries, diabetes, hypertension, and obesity in middle age [10]. Understanding the unique role and interactions of risk factors can allow for early and more personalized interventions, novel targets, and preventive strategies for successful aging.

Pathological changes may be present in the brain decades prior to AD clinical symptoms [11], thus identifying subjects at risk, and early biomarkers can significantly increase the efficacy of interventions. Brain networks, or connectomes, can inform on disease etiology, progression, and response to treatments in humans [12-14], and animal models [15-18]. Connectome properties are preserved across species, providing a translational bridge between preclinical and clinical studies [19, 20]. Understanding the dynamics of the connectome in relation to genetic and other risk factors is of great interest [21, 22], yet most studies have examined one risk factor at a time. While AD is highly heterogeneous, mouse models offer the ability to control both genetics and environment, including exposure to modifiable risk factors. Revealing their impact on connectomes may identify vulnerable regions and circuits, to help disentangle disease heterogeneity [23].

Mice provide tools to study the interaction of AD risk factors along the life span, since they age faster than humans; and replicate the main functional networks in the human brain, e.g. the default mode network [24]. To reveal changes in functional and structural networks before disease onset Chen and colleagues [25] compared elderly APOE4 and non-APOE4 carriers with normal cognition, showing lower global efficiency, and functional connectivity loss in medial temporal areas in APOE4 carriers. The parahippocampal gyrus had functional and structural damage, and its efficiency mediated the APOE4 effect on memory. In mice, APOE4 and APOE-KO genotypes affected functional connectivity independently of age, which was lower for the auditory, motor, somatosensory and hippocampal areas, with APOE-KO accelerating decline in motor, visual and retro splenial cortices [26, 27]. Most research has addressed functional networks, but investigating structural networks and the interplay between these two can provide novel insight. For example, reduced functional and structural coupling has been observed in sensory motor regions during aging, but was preserved for areas of higher cognitive function [28]. LOAD causes aberrant structural connectivity in the prefrontal areas, and temporal pole [21], accompanied by functional changes in the prefrontal, cingulate [29], and temporal cortices [30]. The literature is less clear on the ability to detect vulnerable networks before symptoms onset [31].

Here we examined the impact of fixed and modifiable risk factors, and their interactions, revealing how they converge onto on a small number of vulnerable networks, in mouse models of aging, expressing human APOE alleles. We also evaluated the impact of humanized innate immunity, by replacing the mouse mNos2 gene with the human NOS2 gene, reducing the levels of nitric oxide (NO) production to provide a more similar redox activity to humans [32]. We sought to understand how APOE interacts with age, sex, diet and immunity to confer vulnerability to brain networks estimated from diffusion MRI. MRI was accelerated 8 times by the use of compressed sensing, allowing us to reconstruct diffusion images at 45 μm resolution, segment 332 brain regions [33] [34], and reconstruct connections based on tractography. Connectomes were tested using graph based approaches to estimate risk associated network fingerprints [35], and provide confidence bounds [36]. **Figure 1A** shows a schematic of our approach to reveal networks vulnerable to unique and multiple interacting LOAD risk factors.

**Figure 1.**
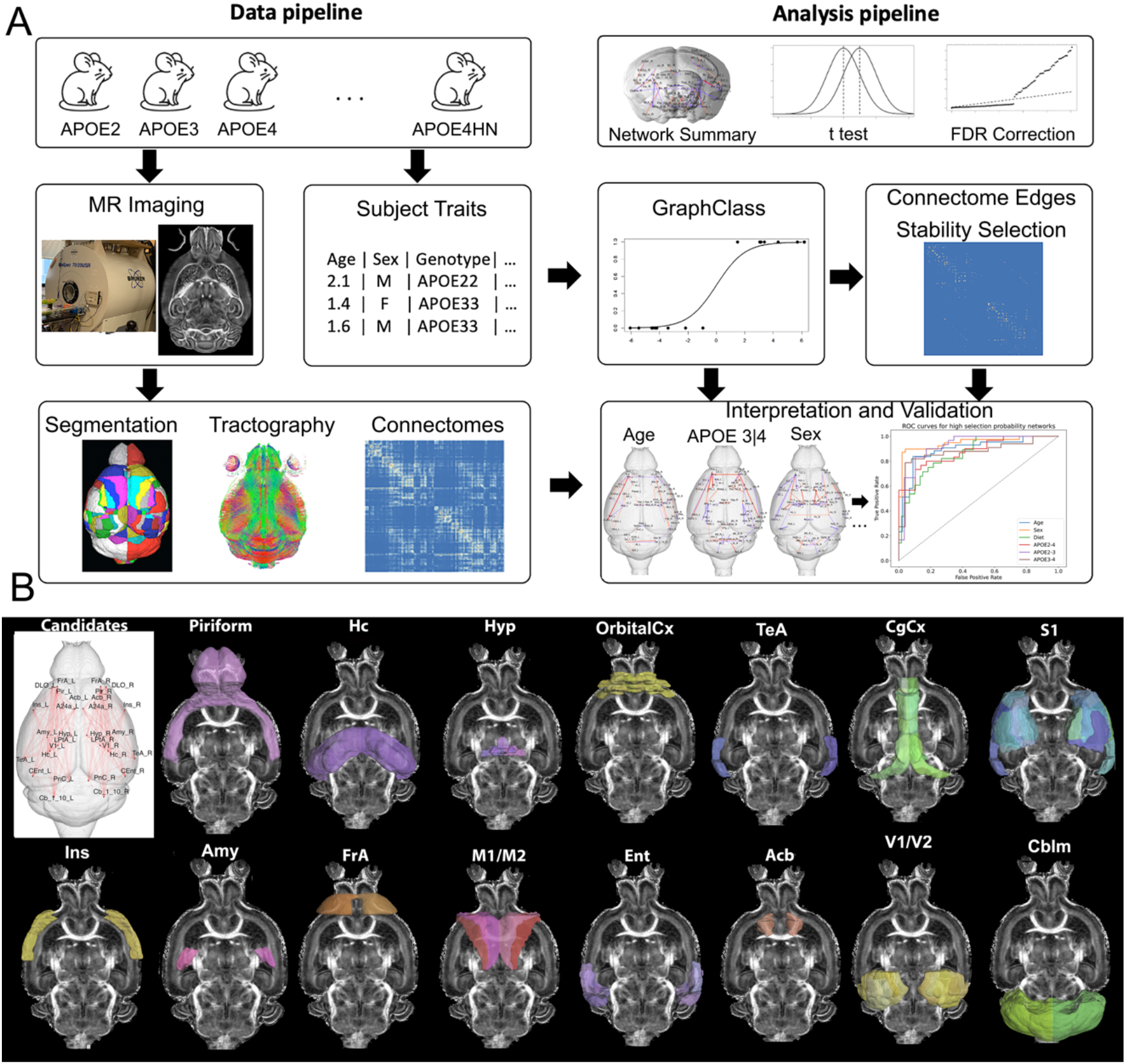
The approach for detecting networks impacted by distinct AD risk factors (APOE genotype, age, immune factors, sex, and diet) comprises a data preprocessing and network analysis pipeline with model validation. We leveraged mouse models expressing various combinations of risk traits, imaged using compressed sensing diffusion weighted MRI. Brains were parcellated using atlas-based segmentation, which was combined with tractography to construct connnectomes; every entry in the connectome matrix represents the number of streamlines that connect each region pair. The analysis pipeline used the network summary statistics in an initial exploratory analysis, followed by graph class and validation of the resulting networks (A). The candidate networks hypothesized to change when varying the levels of risk traits included: the piriform cortex (Pir), hippocampus (Hc), hypothalamus (Hyp), orbital cortex (OrbitalCx), temporal association cortex (TeA), insula (Ins), amygdala (Amy), frontal association cortex (FrA), motor (M1, M2) and entorhinal cortex (Ent), accumbens (Acb), cingulate cortex (Cg), primary somatosensory cortex (S1), visual cortex (V1/V2), and cerebellum (Cblm).

We hypothesized that module hubs for the default mode and lateral cortical networks [37], including the anterior cingulate and frontal association cortices, are also candidates for structural networks. Candidate hubs include the dorsal hippocampus (hippocampus module), accumbens and olfactory nuclei (basal ganglia), pons/ventral subiculum (ventral midbrain), and centromedial thalamic nuclei (thalamus module). The temporal association cortex, cerebellar nuclei and pons are also candidates, since they serve as connector hubs, with high connectivity between modules [38]. **Figure1 B** illustrates network regions hypothesized to be susceptible to fixed and modifiable risk factors. Our study aims to reveal networks vulnerable to specific risk factors, and shared networks vulnerable to multiple risk factors. Such information may be valuable in providing sensitive circuit targets for preventive or pharmaceutical interventions.

## Results

To understand how risk factors impact connectomes to increase susceptibility to LOAD, we first examined global effects through topological network parameters, followed by local impact to determine the network fingerprints for each distinct risk factor, and finally determined common vulnerable networks shared by all risk traits.

### Group-level Differences in Network Parameters Differentiate with LOAD Risk Factors

Exploratory analysis revealed global differences for key network parameters, e.g. eigenvector centrality, betweenness centrality, global efficiency, local efficiency, average clustering, shortest path length, and degree Pearson correlation (**Table 1)**. Aging (comparing above and below median age) was associated with significant differences in eigenvector/betweenness centrality, local efficiency, average clustering, and shortest path length, suggesting extensive differences in connectivity, information flow, and resilience. No significant effect was found for different diets. Local efficiency and shortest path length varied with sex, suggesting innate differences in resilience and information flow. All measures except local efficiency and degree Pearson correlation were significantly different when altering innate immunity (comparing mNOS and hNOS), indicating strong differences in connectivity, information flow, and resilience. All pairwise APOE allelic comparisons had significant differences in eigenvector and/or betweenness centrality, suggesting strong differences in connectivity. Average clustering was significant only for APOE2/APOE3. The degree Pearson coefficient was only significant for APOE genotypes, i.e. for APOE2/APOE3, and APOE2/APOE4, highlighting variations in network assortativity for the APOE2 genotype.

**Table 1.** FDR corrected p-values for t-tests comparing topological summary statistics. (*p<0.05, **p<0.005).

|  | Age | Diet | Sex | mNos/hNOS | APOE2-3 | APOE2-4 | APOE3-4 |
| --- | --- | --- | --- | --- | --- | --- | --- |
| Eigenvector centrality | 0.0017** | 0.78 | 0.58 | 0.039* | 1.8x10<sup>-5**</sup> | 0.29 | 0.03* |
| Betweenness centrality | 0.78 | 0.073 | 0.45 | 0.017* | 2.9x10<sup>-7**</sup> | 3.5x10<sup>-5**</sup> | 0.073 |
| Global efficiency | 0.20 | 0.69 | 0.38 | 0.0063* | 0.24 | 0.75 | 0.36 |
| Local efficiency | 0.044* | 0.79 | 0.022* | 0.38 | 0.20 | 0.72 | 0.36 |
| Average clustering | 0.0037** | 0.10 | 0.31 | 2.0x10<sup>-6**</sup> | 0.039* | 0.75 | 0.20 |
| Degree | 0.20 | 0.66 | 0.36 | 0.0056* | 0.24 | 0.75 | 0.36 |
| Shortest path length | 0.0053* | 0.40 | 0.022* | 0.022* | 0.9 | 0.24 | 0.14 |
| Degree Pearson correlation | 0.72 | 0.78 | 0.66 | 0.91 | 1.45x10<sup>-5**</sup> | 0.0009** | 0.38 |

### Vulnerable Networks Associated with Individual LOAD Risk Factors

We identified subgraphs associated with risk factors such as APOE genotype, age, sex, diet, and immune background (**Figure 2, Dataset S1**), discriminating amongst mouse models with different levels of each trait. To aid with interpretability, we retained the highest ranked 50 pairwise connections (edges) within each high selection probability (HSP) subnetwork, noting that the same false selection bounds still apply to this subpopulation of edges.

**Figure 2.**
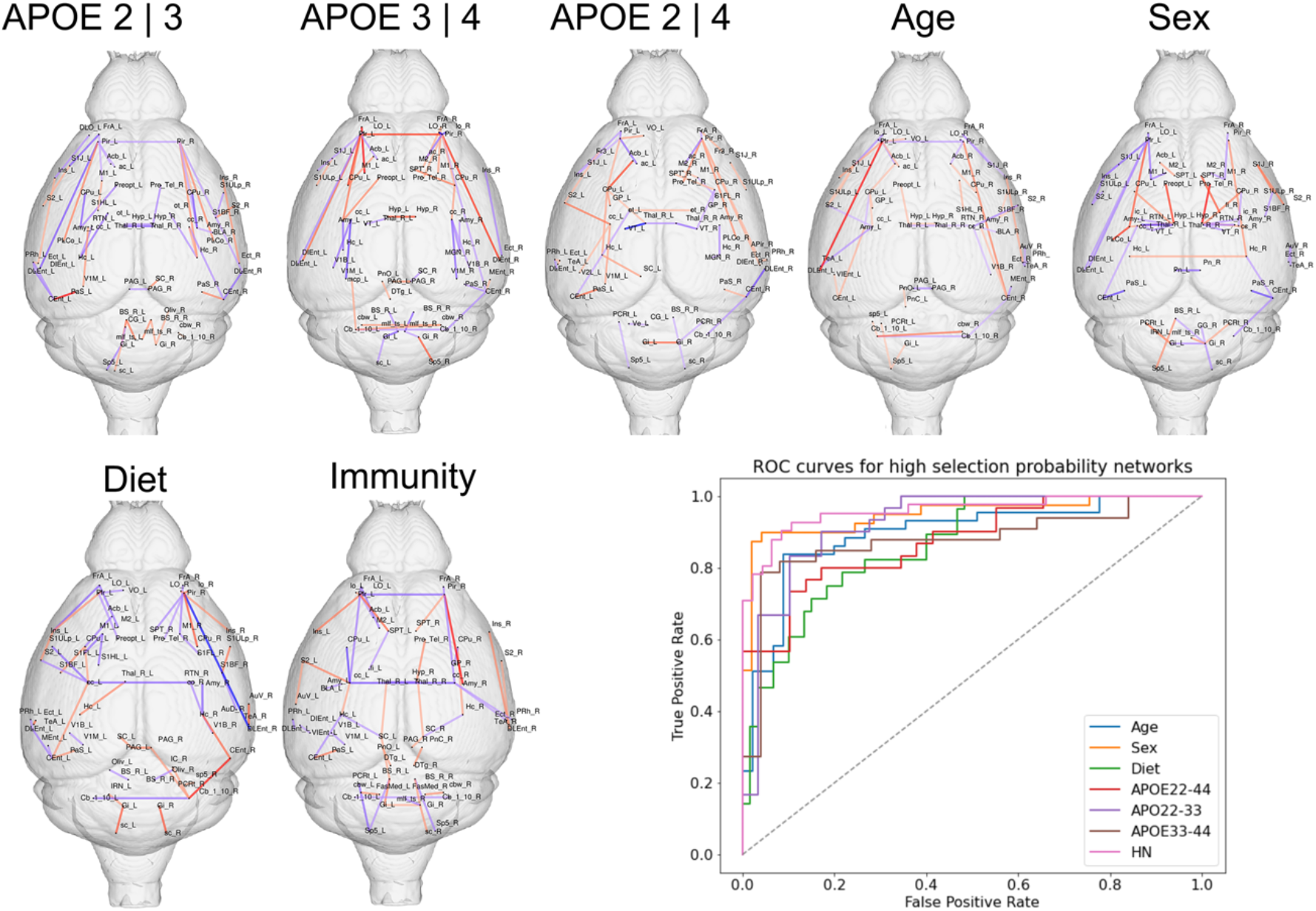
Network fingerprints associated with APOE genotype, age, sex, diet and immunity, and their validation ROC curves. We noted a central role for the entorhinal cortex, parasubiculum, dorsolateral orbital cortex and insula for the APOE2/APOE3 comparison; the frontal association cortex, M1, and a large network including limbic regions, striatum and piriform cortex for APOE3/APOE4; the frontal association cortex, entorhinal cortex, parasubiculum, basal ganglia, insula for APOE2/APOE4. Age also impacted circuits including the entorhinal and piriform cortex, and also giganto cellular reticular nuclei. Sex differences impacted the hypothalamus and amygdala, and also the hippocampus, septum, and fimbria - linking sex vulnerability to memory function. Diet impacted the hippocampus, amygdala and entorhinal cortex, insula, accumbens, and olfactory areas. Immunity (HN presence) showed a role for the frontal association cortex, hypothalamus, piriform cortex and amygdala, and interhemispheric connections though the corpus callosum. Red denotes positive, blue denotes negative edge weights.

#### APOE2/APOE3

We hypothesized AD vulnerability increases from APOE2 to APOE3 to APOE 4. Comparing the neuroprotective APOE2 allele to the neutral APOE3 resulted in 273 connections or edges, with <8 false selections. The most significant subnetwork (BIC 107.4) included 7 regions: the dorsolateral orbital cortex, caudomedial and dorsolateral entorhinal cortex, insula, ectorhinal cortex, parasubiculum, and perirhinal cortex. Interestingly, weights were higher for the left hemisphere, yet contralateral nodes were also represented in the subgraph. The next ranked subnetwork (BIC=126.5) included 16 regions, i.e. the entorhinal cortex (dorsolateral and intermediate), bilateral piriform cortex and amygdala (posterior cortical amygdaloid nucleus, basolateral, rest of amygdala), hippocampus, parasubiculum, accumbens and striatum; and the white matter of the anterior commissure (Left, or L), corpus callosum (Right, or R), and optic tracts (R). The 3^rd^ ranked network (BIC=131.3) included the preoptic telencephalon, striatum and amygdala. Alterations were observed in subnetworks of the limbic system, and in brain regions related to executive function, memory and sleep, as well as in interhemispheric connections.

#### APOE3/APOE4

Since APOE4 is the most significant genetic risk for AD, we compared APOE4 carriers relative to the control APOE3 allele, and identified a vulnerable graph comprising 773 edges with <40 false selections. Massive networks with looser false selection bounds were observed. The two highest ranked networks included connections between left frontal association cortex and primary motor cortex (M1) (BIC=147.5); and between visual cortex and corpus callosum (BIC=148.8). Subsequent ranked subnetworks involved the brain stem, giganotocellular nucleus, trigeminal and tectospinal tracts, the periaqueductal gray and superior colliculus. A large subgraph (33 regions) included the insula, lateral orbital cortex, entorhinal and piriform cortex, parasubiculum, hippocampus, striatum, amygdala, accumbens, and cerebellum, as well as V1, medial geniculate and cerebellar peduncle, preoptic telencephalon and septum (BIC=156.9). Thus, in addition to executive function and memory networks, observed when comparing APOE2 versus APOE3, a sensorimotor component was revealed between APOE3 and APOE4, as a potential biomarker.

#### APOE2/APOE4

We compared the protective versus the high risk APOE allele and identified 204 edges with <4 false selections, the sparsest subgraph of the pairwise APOE genotype comparisons. The highest ranked subnetwork comprised 21 regions (BIC=104.8), including the frontal association cortex, insula, ventral orbital cortex, entorhinal cortex, piriform cortex, septum, parasubiculum and hippocampus, ventral thalamus, striatum, globus pallidus, accumbens, and superior colliculus. The 2^nd^ subnetwork included the frontal association cortex, frontal cortex area 3, the entorhinal cortex, piriform cortex, amygdala, and amygdala-piriform transition area, preoptic telencephalon and anterior commissure. The third network included the primary somatosensory cortex (S1, forelimb area), M1, and M2. Subnetworks involved in executive function, memory, as well as sensorimotor areas, sleep, and reward processes, were identified in this fingerprint.

#### Age

Since age is the largest risk for AD, we compared animals below and above the median age of 16 months, which corresponds to late middle age, and obtained 300 high selection edges with <5 false selections. The top 50 edges included interhemispheric periaqueductal gray connections, involved in responses to threatening stimuli (BIC=211.2); the spinal trigeminal and gigantocellular reticular nuclei (BIC=213.8); the temporal and frontal association cortices, S1, secondary auditory cortex, ectorhinal and perirhinal cortex (BIC=239.4). Thus, age impacted circuits involved in executive, sensory and motor functions, and in the response to stressors. We also observed changes to subnetworks including the hypothalamus, S1, S2, V1, and the pons. The largest high selection probability (HSP) subnetwork included 31 regions, e.g. the frontal and temporal association cortex, ventral orbital cortex, S1, S2, entorhinal and piriform cortices, amygdala, accumbens, globus pallidus and striatum, as well as the cerebellum gray, and white matter. Other connections involved the anterior commissure, lateral olfactory tract, trigeminal nerve, and corpus callosum. Unsurprisingly, age influenced a large portion of the brain connectome.

#### Sex

Because females constitute 2/3 of AD patients, we compared female to male connectomes, and obtained 540 high selection probability edges with <7 false selections. The top subgraph (BIC=201.4) included the reticular nuclei (parvicellular, principal sensory trigeminal, intermediate), gigantocellular, medial longitudinal fasciculus, tectospinal tract. The 2^nd^ and 3^rd^ subgraphs (BIC=203.2 and 213.5, respectively) included the right (2^nd^ subgraph) and left (3^rd^ subgraph) thalamus, ventral and reticular nuclei, internal capsule, and striatum. The corpus callosum was also part of the 2^nd^ subgraph. The 4^th^ network (BIC=215.5) included the entorhinal cortex, parasubiculum, hippocampus and cerebellum; and the 8^th^ included the preoptic telencephalon, amygdala and hypothalamus. The hypothalamus influences energy balance and metabolism, and the gigantocellular component supports differences in recovery after injury [39].

#### Diet

Since diet can significantly impact brain milieu, e.g. providing nutrients, modulating glucose metabolism, oxidative stress, inflammation, and insulin signaling, we compared the impact of a high fat versus regular diet and identified 1626 high selection probability edges with <31 false selections. The top subgraph (BIC=193.5) included the caudomedial entorhinal cortex, hippocampus, amygdala, reticular nucleus of thalamus, periaqueductal gray, superior colliculus, cerebellar cortex, and primary visual cortex (V1). The 2^nd^ subgraph (BIC=200.25) included the insula, piriform, lateral orbital and dorsolateral entorhinal cortex, septum, preoptic telencephalon, striatum and the lateral olfactory tract. The 3^rd^ and 4^th^ graphs related the olivary complex with the brain stem rest (BIC=201.5, 204.5). The auditory and temporal association cortices were also present in top subnetworks. Diet impacted circuits involved in sensory processing, reward and motor control, emotion regulation, response to stress and inflammation, and memory.

#### mNos/hNOS

Since immune alterations have been associated with AD, we performed a comparison of high versus lower NOS production (moving from mNos to hNOS) and identified 411 edges with <9 false selections. The top subgraph (BIC=165.3) was composed of 32 regions, including the secondary auditory cortex, somatosensory cortex and motor cortex, as well as the frontal association cortex, perirhinal and entorhinal cortices, parasubiculum, hippocampus, hypothalamus, thalamus, superior colliculus, striatum, globus pallidus, preoptic telencephalon, gigantocellular reticular nuclei, periaqueductal gray, areas of the brain stem, plus tracts i.e. the corpus callosum, spinocerebellar, tectospinal, medial longitudinal fasciculus. The 2^nd^ subgraph (BIC=171.8) included the piriform cortex, septum, accumbens, amygdala, fimbria, and the olfactory tract. Interestingly the 3^rd^ and 4^th^ networks included the cerebellar fastigial nucleus and white matter. The insula and entorhinal cortex were found in the next ranked subgraph (BIC=216.8). The pontine reticular nuclei and dorsal tegmentum were also present in the top ranked subnetworks. Thus, we identified extensive networks that interact with immune mechanisms, influencing and being influenced by neuroinflammation and immune responses.

### Validation of Vulnerable Networks Associated with Individual Risk Factors for LOAD

High selection probability networks were validated by predicting each trait using only edges in the selected network. The areas under the curve (AUC) across 10 train split tests were: 0.96±0.03 for sex; 0.95±0.02 for immunity (mNos/hNOS), 0.90±0.03 for age, 0.860±02 for diet; 0.92±0.03 for APOE2 vs APOE3, 0.87±0.03 for APOE2 vs APOE4, 0.87±0.04 for APOE3 vs APOE4 (**Figure 2**). Thus, all models performed very well, despite the extreme sparsity of the selected networks

### Brain Networks with Shared Vulnerability Across Risk Factors for LOAD

To reveal vulnerable networks associated with multiple risk factors, we pooled the independent analyses and retained the top 50 edges ranked by cumulative weight across all the HSP networks (**Figure 3A and C, Dataset S2)**.

**Figure 3.**
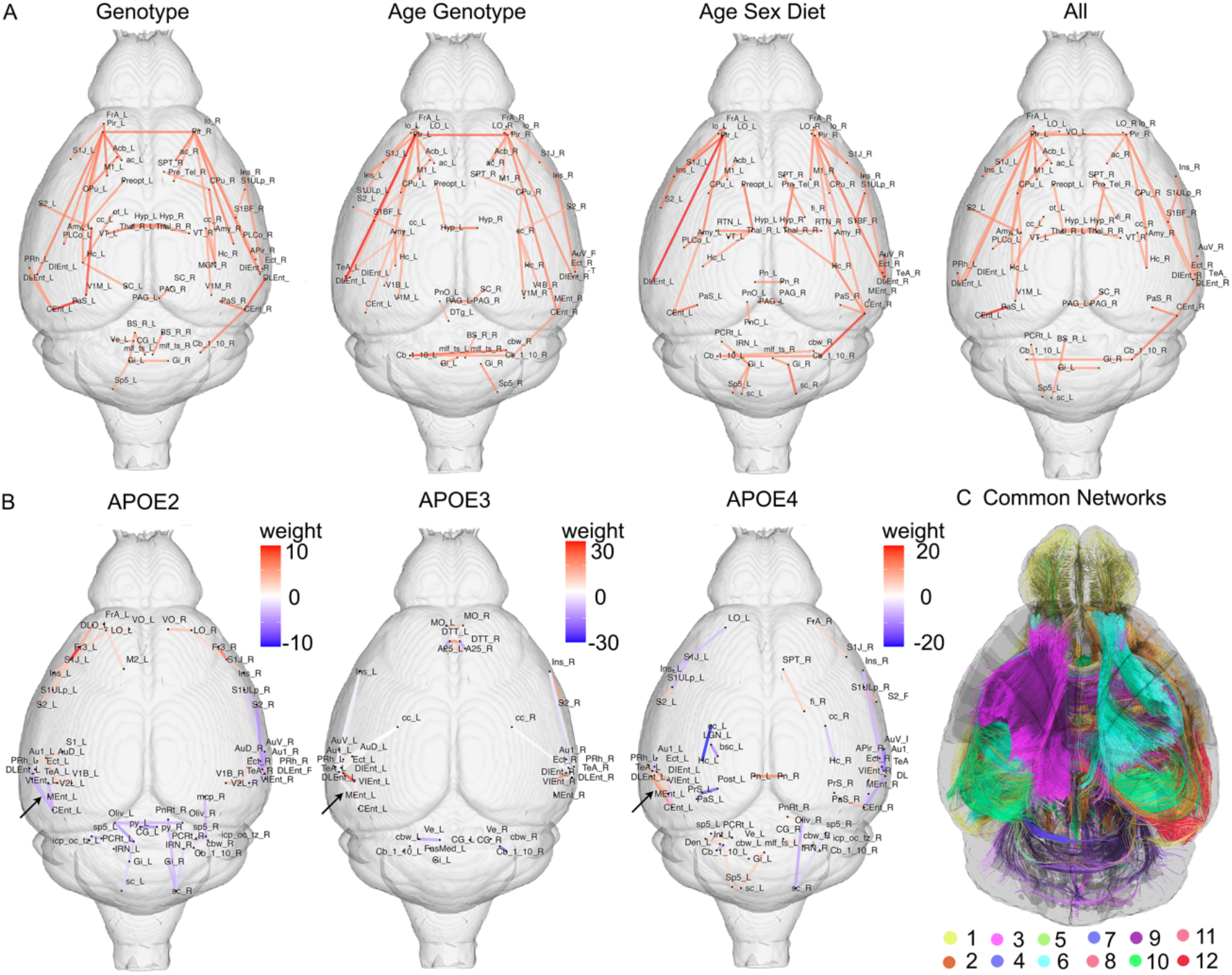
Common edges for vulnerable networks associated with multiple risk factors for. LOAD, and their interactions (A). The immune changes due to HN affected similar networks in different APOE genotypes, including memory networks, with a strong entorhinal circuitry component, but also S2, the auditory cortex, insula and cerebellum. Relative to the neutral APOE3 allele, the connections within the entorhinal cortex (black arrows) in APOE2 mice had negative weights, while in APOE4 mice they had positive weights (Caudal to Dorsal Entorhinal cortex). Common networks with shared vulnerability for APOE genotype, age, sex, diet and immunity (C).

#### APOE Genotype

110 edges were sensitive to variation in APOE genotype for all paired comparisons. The largest subnetwork involved 25 regions, including the bilateral dorsal intermediate entorhinal cortex, posterolateral amygdaloid area, piriform cortex, hippocampus, amygdala, striatum, preoptic telencephalon, and anterior commissure. The visual cortex, accumbens, superior colliculus, medial geniculate and septum were also part of this large subnetwork. Interhemispheric tracts e.g. the corpus callosum, optic and olfactory tracts emerged as important. The 2^nd^ network involved the caudomedial and dorsolateral entorhinal cortex, parasubiculum, and perirhinal cortex. The 3^rd^ network included the contralateral components for the first 3 regions and the cerebellar cortex. Among the remaining subnetworks we noted the frontal association cortex, interhemispheric hypothalamic and ventral thalamic connections, S1, and S2, M1, insula, ectorhinal cortex, and periaqueductal gray. The gigantocellular reticular nuclei and tectospinal tract were amongst the top 13 subnetworks. Thus, the APOE sensitive networks included regions vulnerable in AD, and a set of regions with roles in integrating information and orchestrating cognitive, emotional, behavioral responses, sensory and motor control.

#### Age and APOE Genotype

239 edges were common for the risk factors represented by age and APOE4 versus
APOE3 allele carriage. The top subnetwork involved 24 regions, 6 unique to the left hemisphere, and 2 unique to the right hemisphere. The 8 nodes with bilateral presence included the lateral orbital cortex, the dorsal intermediate and caudomedial entorhinal cortex, piriform, accumbens, cerebellar cortex, lateral olfactory tract and anterior commissure. The nodes present only in the left hemisphere included the dorsolateral entorhinal cortex, S1 (barrel field), insula, amygdala, hippocampus, striatum; the nodes only present in the right hemisphere included the medial entorhinal cortex, and cerebellar white matter. The 2^nd^ subnetwork included the right septum, hippocampus, striatum, S2, V1, and corpus callosum. The 3^rd^ subnetwork included the frontal association cortex, insula, S1, M1, ectorhinal cortex. We noted the presence of the temporal association cortex, hypothalamus and preoptic telencephalon, visual and auditory cortices, periaqueductal gray, gigantocellular nucleus, tectospinal tract. While we recognized the presence of top subnetworks including regions responsible for APOE genotype differences, we also noted an additional cerebellar component.

#### Age, Sex, and Diet

197 edges were common for the risk factors represented by age, sex and diet. The top larges subnetwork comprised 19 regions, including the bilateral cerebellar cortex, right hippocampus, entorhinal cortex, amygdala, piriform, parasubiculum, amygdala, lateral olfactory tract, right striatum, reticular thalamic nuclei, temporal association cortex, insula, and auditory cortex. The 2^nd^ subnetwork included contralateral components, i.e. left entorhinal cortex, amygdala, piriform, parasubiculum, amygdala, lateral olfactory tract. The 3^rd^ subnetwork included 6 regions in the left hemisphere, i.e. the frontal association cortex, S1, S2, M1, lateral orbital cortex, and insula. We noted the hypothalamus, septum and fimbria, periaqueductal gray, gigantocellular nuclei, and contralateral components for regions in the top subnetworks. Thus memory, sensory, emotion, and motor circuits, were identified as networks with shared vulnerability to these three risk factors.

### APOE Genotype Specific Vulnerability to Immune Perturbations

We examined APOE genotype specific connectome fingerprints associated with the manipulation of the innate immune system. For the APOE2 versus APOE2HN comparison we obtained 265 HSP with upper bound of 185 false discoveries; for APOE3 versus APOE3HN we obtained 47 HSP with less than 3 false discoveries; for APOE4 versus APOE4HN we obtained 1091 HSP with less than 598 false discoveries. Larger networks and numbers of regions were found in APOE4 (61 regions) relative to APOE2 (39 regions) and APOE3 mice (40 regions). The innate immune changes affected memory networks, and regions changing early in AD for all genotypes, such as the entorhinal and temporal association cortex; as well as the ventral orbital, ectorhinal and perirhinal cortex, secondary somatosensory cortex (S2) and gigantocellular nucleus, and the auditory cortex. However, APOE3 had almost neutral effects for connections within the entorhinal cortex, APOE2 mice had negative and APOE4 mice had positive weights associated with connections within the entorhinal cortex (black arrows in **Figure 3B**) between Caudal to Dorsal Entorhinal cortex). Upon examining the top 50 edges we noted that APOE2 mice spared the corpus callosum, APOE3 mice had changes in the cingulate cortex and corpus callosum, and APOE4 mice had changes in the callosum-hippocampus connectivity, bilaterally, but with a stronger weight for the left hemisphere.

### Common Edges

63 edges, constituting 12 subnetworks, were common for age, sex, APOE genotype, HN genotype, and diet (**Figure 3C** shows the top subnetworks, ranked by weight). The top subnetwork included 29 regions, e.g. bilateral entorhinal cortex, hippocampus, amygdala, striatum, piriform cortex, and the anterior commissure and corpus callosum; the left ventral orbital cortex, V1, S2, accumbens, perirhinal cortex; and right parasubiculum and lateral orbital cortex. The 2nd subnetwork included interhemispheric thalamic connections. The 3^rd^ network included hypothalamus, preoptic telencephalon connections. The other subgraphs included frontal association cortex, insula, S1 and M, septum, fimbria, temporal association areas, auditory cortex, fimbria, olfactory and optic tracts. The periaqueductal gray and superior colliculus were also present, as well as gigantocellular reticular nucleus. Thus, a small set of edges (0.11 % of the total 54946 edges in the mouse brain connectomes) were common to all the risk factors.

## Discussion

Our study leveraged mouse models to reveal network fingerprints associated with LOAD risk factors i.e. APOE genotype, age, sex, and innate immunity. The most frequent differences (3 of the 6 comparisons) were observed for eigenvector centrality describing the influence of nodes on a network, betweenness centrality describing the importance of a node in controlling the information flow within a network, clustering coefficient which measures the degree to which neighboring nodes aggregate, i.e. local connectivity, and shortest path length related to global integration. Two comparisons were significant for local efficiency (the inverse of the shortest path length for all neighbors of a node), and assortativity. Our results support that in addition to age, immunity strongly impacts connectivity, information flow, and network resilience; and APOE and sex impact networks architecture in the absence of AD pathology.

We obtained mainly symmetrical networks, although weights appeared generally higher for the left hemisphere. When comparing APOE genotypes, the largest networks, and highest number of selection probability edges were detected when comparing APOE3 to APOE4 mice, i.e. 1.4 % of the total number of connections or edges in a connectome, followed by 0.5% for APOE2 versus APOE3, and 0.4% for APOE2 versus APOE4. Surprisingly, APOE2 connectomes were more similar to APOE4 connectomes, which is in agreement with a similarity in spatial navigation strategies of APOE2 and APOE4 mice [40]. APOE2 versus APOE3 differences involved the lateral orbital cortex, entorhinal and perirhinal cortex. Besides regions involved in executive function and memory, the APOE3 versus APOE4 comparison emphasized the role of sensory and motor areas including vision, taste, and pain (insula, periaqueductal gray). Sensory and motor associated regions differed in asymptomatic APOE4 carriers relative to APOE3 carriers [41]. Here we also observed connectivity differences for the cerebellum, well-known for its role in movement control, and emerging as important in language, learning and memory function [42]. Our results also support a significant role for the basal ganglia and amygdala connectivity, involved in affective memory.

We identified the largest networks for diet, followed by sex, immunity, and age. The large number of impacted edges for diet (1626) is likely the consequence of it being an intervention, whereas the low number for age (∼300) reflects healthy aging. Similar to humans [43], mouse networks changed with aging, which impacted association, motor and sensory circuits, e.g. olfactory, the basal ganglia and cerebellum.

Our results help understanding sex specific vulnerability [44], as this network fingerprint pointed to sexually dimorphic regions i.e. the basal ganglia, hypothalamus (48[45], hippocampus, and amygdala [46]. Memory related circuits (including the entorhinal cortex, hippocampus, septum and fimbria [47] [48], were supplemented with cerebellum and gigantocellular nuclei circuits, suggesting differences in locomotor recovery after injury[39] [49].

Diet impacted networks involved in appetite control, e.g. insula and hypothalamus [50] [51] [52], which was also impacted by immune differences. The hypothalamus triggers inflammatory responses following a high-fat diet, which have downstream effects on synaptic plasticity, neurogenesis and neuromodulation [53]. The finding of visual, auditory and cerebellar regions corroborate human studies showing changes in somatomotor, and visual areas, and suggest diet impacts memory (entorhinal cortex, and hippocampus) [54], and attention (parahippocampal gyrus) networks, possibly accelerating brain aging [55].

The immune network fingerprint emphasized a role for reticular nuclei, which may act either through autonomic modulation, or indirectly on the hypothalamus. The corpus callosum, piriform cortex and amygdala were impacted across all APOE genotypes. Within genotype comparisons consistently converged onto memory and sensory networks, including S2, and the auditory cortex. Significantly larger networks were observed for APOE4, relative to APOE2 and APOE3, suggesting increased sensitivity to immune perturbations for APOE4 genotypes. The interactions of APOE with aging impacted large limbic circuits (hippocampus, entorhinal cortex, amygdala), interacting with sensory and motor areas, the reward system (insula, accumbens, lateral orbital cortex, striatum), association cortices; and impacted anterior commissure and olfactory tract circuitry. Our results parallel human studies showing that APOE4 carriers had accelerated structural connectivity loss in the inferior temporal, medial and lateral orbital cortex [56], and support that olfaction may provide early markers of risk for accelerated aging, and the need for further probing the specific role of the highly connected cerebellum in AD [57].

Common networks for age, sex and diet included S1, S2, M1, lateral orbital and entorhinal cortex, insula, septum, amygdala and hypothalamus, fimbria and olfactory tract, in agreement with aging human functional connectomes, and faster cortical thinning in aging, e.g. the temporal and sensory-motor areas. Our study identified the auditory cortex, while the human study identified the visual cortex, however this only accounted for aging but not sex and diet associated vulnerability.

The network with shared vulnerability for all risk factors consisted of only 63 edges, and 12 subnetworks, and supported a role for the orbitofrontal cortex, association cortices, insula, S1, M1, auditory cortex, and cerebellum in addition to well-known AD vulnerable regions, and their connections. These results underline the importance of early sensory motor markers in AD.

Most preclinical MRI studies have examined functional brain networks in models of familial AD, thus in the presence of AD pathology. The 5xFAD mouse has lower clustering coefficients, small worldness, and modularity in functional networks including: the hypothalamus, superior colliculi; amygdala, brainstem, central gray, cerebellum, pallidum, hippocampus, inferior colliculus, neocortex, olfactory, midbrain, basal forebrain/septum, thalamus [19]. In models of genetic risk for LOAD, APOE4 as well as APOE-KO genotypes had different functional connectivity independently of age, in particular for the auditory, motor, somatosensory and hippocampal area, and APOE-KO mice had faster decline in motor, visual and retro splenial cortices [26, 27]. How AD risk impacts structural mouse brain networks has been less studied, but the strong link between structural and functional connectomes led to many of the regions detected by our structural analyses to appear as critical for predicting functional alterations in networks containing these nodes.

Our study adds a structural, fine grained anatomical substrate to support previously reported alterations of mouse functional networks, including the default mode network, the lateral cortical network, hippocampus, basal forebrain, ventral midbrain, and thalamus [58]. While it has been shown that aging differentially affected these networks [38], the effect of multiple risk factors has been insufficiently addressed. We revealed alterations in structures involved in the default mode network, salience network and executive control network that are likely to impact memory retrieval, and executive function in the absence of pathology, in models of preclinical AD. This supports the importance of preclinical human studies in populations at risk. Our findings of impacted sensory networks suggest potential early biomarkers based on auditory, olfactory, and sensory motor tests.

Targeted replacement APOE mice have differences in behavior, cerebral perfusion and functional connectivity [59], [60] [3] [61]. The hNOS2 gene has been linked to oxidative processes, DNA repair, and mitochondrial activity, altering TNFα and CCR1 mRNA expression [62, 63]; our study shows that immune changes affect brain networks in an APOE dependent fashion.

Our study has limitations since mouse models do not fully replicate the complexity of human AD, however they provide a uniform genetic background and control over risk factors; enabling smaller studies to reveal mechanistic insight into disease etiology, and testing the ability to curb its evolution, through interventions such as diet or exercise, in a causal connectomic approach.

Our method provides stable network estimates with bounds for false discoveries, carefully controlling for multiple comparisons. The false discovery rate bounds hold under general conditions, and remain valid for subsets of selected edges. The comparison of the APOE4 high-risk genotype, to the neutral APOE3 genotype revealed vulnerable networks; while that of the protective APOE2 to the neutral APOE3 revealed potentially resilient networks. We showed that immunity impacted memory and sensory networks, but future studies including more animals shall further assess genotype specific differences. We identified most candidate regions hypothesized to change with AD risk, with the exception of the cingulate cortex, only present in one of the comparisons. The small size of cingulate cortex subdivisions likely influenced our ability to detect changes, and this could be improved by using a hierarchical parcellation scheme.

Our results revealed network fingerprints for unique AD risk factors, and provided insight into networks with shared vulnerabilities to multiple risk factors. These networks may provide the substrates underlying functional network alterations, but at higher anatomical granularity than fMRI. Further mechanistic studies may reveal metabolic and omic bases for structural networks. Our results provide motivation for exploring structural networks as early neurodegeneration markers, to enable designing more targeted interventions to delay onset or progression of AD.

## Materials and Methods

### Animals

Animal procedures were approved by the Duke Institutional Animal Care and Use Committee. To model genetic risk for LOAD, we used mice lacking the mouse Apoe gene, and homozygous for the three major human APOE alleles [64, 65]: the high-risk gene APOE4, the control APOE3 and the protective APOE2 allele. These lines were crossed with mice where the mouse Nos2 gene has been replaced with the human NOS2 gene (HuNOS2tg/mNos2^-/-^ mice, termed HN) [62, 63]. This modification addresses, in part, the differences between the human and mouse inflammatory responses, where human, in contrast to mouse macrophages, express little NOS2, and generate much less NO in response to inflammatory stimuli [66]. Introducing the human NOS2 gene changes redox balances to better mimic those in the human brain. Removing the mNos2 gene, and introducing the human NOS2 gene lowers the amounts of NO produced, and brings the mouse immune/redox activity more in tune with the human redox activity, which promotes AD pathologies [62]. Animals were bred as homozygous for each of the APOE alleles, either to completely lack the mNos2 and to express hNOS2, or on a mNos2 background. Both males and females were included, for a total of 173 mice; aged from 13 to 20 months. Of the 59 APOE2 mice, 28/31 were male/female, 15/44 were control/high fat diet, 36/23 were above/below median age, 25/34 were HN/mNOS2; of the 58 APOE3 mice, 28/30 were male/female, 26/32 were control/high fat diet, 27/31 were above/below median age, 23/35 were HN/mNOS2; of the 58 APOE4 mice, 29/29 were male/female, 16/42 were control/high fat diet, 33/25 were above/below median age, and 20/28 were HN/mNOS2.

Thus animals were homozygous for each of the APOE alleles, and either expressing the mNos2 or completely lacking the mNos2 and expressing hNOS2. Both males and females were included, for a total of 173 mice; aged from 13 to 20 months of age, the median age was 16.2 months. These animals were aged naturally, either on a regular chow (2001 Lab Diet) for the whole duration, or switched for 4 months prior to imaging to a high fat diet (D12451i, Research Diets).

These animals were aged naturally, either on a regular chow (2001 Lab Diet) for their whole life duration, or switched for 4 months prior to imaging to a high fat diet (D12451i, Research Diets) containing 45 kcal % fat (39 kcal % from lard; 5 kcal % from oil), 35 kcal % carb (17 kcal % sucrose), and 20 kcal % protein for ∼4 months. Control diet animals received 13.6 kcal % fat, 57.5 kcal carb (3.25 sucrose), and 28.9 kcal % protein throughout their life span. Animals had free access to food and water. The means of each age category were 14.60±1.13 months, and 19.35±1.59 respectively; while the median age was 16.2 months. Animals were sacrificed during a transcardiac perfusion fixation under surgical plane anesthesia with 100 mg/Kg ketamine and 10 mg/Kg xylazine, before being perfused through the left cardiac ventricle, with outflow from the right atrium. Saline (0.9%) was used to flush out the blood, at a rate of 8 ml/min for ∼5 min. For fixation we used a 10% solution of neutral buffered formalin phosphate containing 10% (50 mM) Gadoteridol (ProHance, Bracco Diagnostics Inc., Monroe Township, NJ, United States), at a rate of 8 ml/min for ∼5 min. Gadoteridol reduced the spin lattice relaxation time (T1) of tissue to ∼100 ms, enabling rapid imaging. Mouse heads were trimmed of extraneous tissue, and stored in 10% formalin for 12 h, then transferred to a 0.01 M solution of phosphate buffered saline (PBS) containing 0.5% (2.5 mM) Gadoteridol, at 4°C for ∼30 days to rehydrate the tissue. Specimens were placed in MRI-compatible tubes, immersed in perfluoropolyether (Galden Pro, Solvay, NJ, United States) for susceptibility matching. Specimens were left inside the skull to preserve tissue integrity and shape, but extraneous muscle tissue and the lower jaw were removed to allow close positioning in a tight-fitting solenoid coil.

### Image Acquisition and Processing

Diffusion weighted MRI was done using a 9.4T high field MRI, with a 3D SE sequence with TR/TE: 100 ms/14.2 ms; matrix: 420 × 256 × 256; FOV: 18.9 mm × 11.5 mm × 11.5 mm, 45 μm isotropic resolution, BW 62.5 kHz; using 46 diffusion directions, 2 diffusion shells (23 at 2,000, and 23 at 4,000 s/mm^2^); 5 non-diffusion weighted (b0), as in [40]. The max diffusion pulse amplitude was 130.57 Gauss/cm; duration 4 ms; separation 6 ms, eightfold compressed-sensing acceleration [67] [68]. Diffusion tensor properties such as fractional anisotropy, orientation distribution functions, and tractograms were reconstructed and SIFT filtered using MRtrix3, which was also used for calculating connectomes [69]. We used the SAMBA pipeline implemented in a high-performance computing environment to segment the brain in 332 regions[70], to reconstruct connectomes as adjacency matrices, where each entry represented the number of streamlines connecting a pair of brain regions. The symmetrized mouse brain atlas used by SAMBA to produce anatomical parcellations is available from https://zenodo.org/records/10652239 [68], and the list of brain regions, abbreviations, and indices is provided in **Dataset S3**.

### Exploratory Network Analysis

As a first step we performed an exploratory data analysis to assess global differences for key network parameters in relation to AD risk traits. These parameters included eigenvector centrality, betweenness centrality, global efficiency, local efficiency, average clustering, shortest path length, and degree Pearson correlation, calculated using NetworkX (https://networkx.org/). Eigenvector centrality is a relative score measuring the influence of a node; nodes gain influence by being connected to other influential nodes. Betweenness centrality is a node specific measure counting the number of shortest paths in the graph that pass through a node, and illustrates the nodes control over the network. Global efficiency summarizes the average inverse distance between nodes in a graph. Conversely, local efficiency is a measure of the average inverse distance between a node and its neighbors. The shortest path length between a pair of nodes reflects efficiency of information transfer. Finally, assortativity is Pearson correlation between the degrees of connected nodes (weighted sum of edges), and illustrates the dependency of neighboring nodes. Node specific measures in each brain were averaged to produce mouse specific summaries of connectivity. These measures provided a broad overview of structural variation in connectomes [71]. A t-test was used to detect differences in the summary statistics across risk trait levels, and controlled for false discoveries based on adjusted p-values (FDR =0.05). It is possible to identify subnetworks which vary across groups using node-to-node comparisons, but this approach does not share information across nodes and requires many comparisons, reducing efficiency. In addition, the identified networks are sensitive to the graph metrics used, with no clear optimal choice. For these reasons, we preferred a sparse network classification model.

### Sparse Network Regression

To reveal group differences between APOE2/APOE3, APOE3/APOE4, APOE2/APOE4, mNos/hNOS, below/above median age, female/male, and regular/high-fat diet, we fit a GraphClass [36] sparse logistic regression model to each group. Logistic regression models the probability that a mouse is in one group (e.g., below median age) over another (e.g., above median age) using the vectorized brain connectomes. The probability is obtained by (1) multiplying each edge in the connectome by a coefficient, (2) summing the weighted edges, and (3) transforming this number to be between 0 and 1 so it is a valid probability. Coefficients which best explain the data are estimated via maximum likelihood. Due to the high dimensional nature of the connectomes, it is difficult to interpret the many coefficients. Consequently, we used a sparse model, which penalizes nonzero coefficients and dramatically reduces the number of parameters to visualize.

GraphClass [36] is one implementation of sparse logistic regression which takes advantage of the brain network structure to produce more interpretable results, that is subnetworks that are predictive of each trait. The goal is to estimate the coefficient matrix which best explains the data; by maximizing the likelihood of the observed data. No additional thresholding of coefficients is required after fitting the model.

Our sparse logistic regression models used the GraphClass double sparsity penalty defined in [36]. Let B be a 332×332 symmetric matrix of coefficients, with b_ij_ the coefficient for the edge between regions i and j in a logistic regression model. Let *B*_(*v*)_ be the vth row (or column) of B, v=1,…,332. The GraphClass penalty is:

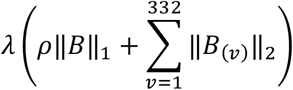

where *λ, ρ* > 0 are sparsity hyperparameters. The first term, ‖*B*‖_1_, is the usual Lasso penalty and shrinks the coefficients of each edge towards zero. The second term, ‖*B*_(*v*)_‖_2_, penalizes the coefficients of all edges connected to region v. In practice, this effectively deletes entire regions from the connectome, resulting in sparser and more interpretable subnetworks. The penalized model is fit with the alternating direction method of multipliers (ADMM). Hyperparameters were selected via 10-fold cross-validation minimizing misclassification rate, and ties were broken by selecting the sparsest model.

### Stability Selection

A subnetwork selected by a single run of GraphClass may be unreliable due to noise in the data and high correlation between entries of the connectomes. Following [36], we used complimentary pairs stability selection (CPSS, [72]) to improve model robustness and reliability. CPSS is based on the intuition that important edges will be selected in a high fraction of repeated experiments (e.g., selecting new mice, reproducing the study, and fitting GraphClass), whereas less important edges will be selected in a low proportion of repeated experiments. The collection of high and low selection probability edges may be estimated via resampling. In CPSS, this entails (1) splitting the data into halves, (2) fitting GraphClass to each half, and (3) saving the selected (nonzero) coefficients from both models. This is repeated, and the proportion of times a coefficient was nonzero across all runs is used to estimate the true unknown selection probability of the corresponding edge. We define high selection probability (HSP) edges as those which appeared in >99% of runs. CPSS comes with theoretical upper bounds on the number of low selection probability edges (LSP) which are falsely labeled as HSP edges; this is a direct analogue of false discovery rate control. We used this procedure with defaults from Shah and Samworth [73] to identify small, reliable subnetworks with false-selection guarantees.

### Subnetwork Ranking

To make the results more interpretable, we ranked the relative importance of connected components within each HSP network by comparing the Bayesian Information Criterion (BIC) of unpenalized logistic regression models predicting the trait using only the edges in a single connected component. BIC balances predictive performance and model complexity, allowing for fair comparisons between subnetworks of different sizes. We consider connected components with lower BIC to be more important for explaining a trait than those with higher BIC.

### Validation

GraphClass and stability selection were used to produce robust subnetworks for predicting each of APOE2/APOE3, APOE3/APOE4, APOE2/APOE4, mNos/hNOS, below/above median age, female/male, and regular/high-fat diet. We validated these subnetworks using AUC/ROC. AUC/ROC was computed by fitting GraphClass regression on half of the samples, using only edges in the HSP subnetwork. This model was used to predict class labels for the other half of the samples. The entire procedure was repeated 10 times with different train/test splits.

### Common Networks

We defined the shared high selection probability (HSP) network across multiple models as the intersection of the HSP networks for each model. For example, the shared networks across age and APOE3/APOE4 contain all the edges that are in the HSP network for age and also in the HSP network for APOE3/APOE4; edges that were only in the HSP network for age or only in the HSP network for APOE3/APOE4 were not included.

## Acknowledgments

We are grateful for NIH support through RF1 AG057895, R01 AG066184, U24 CA220245, RF1 AG070149, and for NSERC support to SW with PGS D578055. We thank Gary Cofer, Wyatt Austin, and Chris Petty for technical support, and Stephen Lisberger for helpful comments.

## Institutional Approval

All animal studies were approved by the Duke Institutional Animal Care and Use Committee.

## Data Sharing Plans

The connectome and associated metadata are available at https://zenodo.org/record/8377684. The software code is shared at https://github.com/szwinter/MouseModels/, and we used visualization code from https://github.com/Ali-Mahzarnia/brainconn2.

## Supplementary Material

**Dataset S1**. Network fingerprints for individual LOAD risk factors

**Dataset S2**. Common Networks

**Dataset S3**. SAMBA Atlas Abbreviations

